# A new reference-invariant consensus template generation method in ALPACA

**DOI:** 10.64898/2026.05.29.728799

**Authors:** A. Murat Maga

## Abstract

1. Automated landmarking transfers anatomical landmarks from a reference specimen onto many targets, greatly increasing analytical throughput. However, this procedure needs to be bootstrap using an initial sample. An arbitrary or atypical choice imprints a reference-of-origin bias that propagates through the pseudo-landmarks, the resulting morphospace, and the downstream template selection, a risk that is difficult to avoid for large datasets whose variation is not yet understood.
2. We replace the fixed reference with an iterative consensus atlas, warped over a few iterations toward the Procrustes mean shape of all similarity-aligned specimens. We evaluated it on a 62-strain *Mus musculus* skull panel by running both the original fixed-reference pipeline and the new consensus pipeline 62 times each, using every specimen in turn as the bootstrap. We compared atlas convergence, inter-atlas similarity, morphospace reproducibility, reference-choice variance of pairwise Procrustes distances, downstream k-means selection stability, and leave-one-out out-of-sample fit, and tested generalisation on great-ape datasets of differing sampling balance.
3. The consensus atlas converged within a few iterations and was far less sensitive to the starting specimen than the fixed reference. It produced more reproducible morphospaces (mean RV 0.960 versus 0.944), reduced the reference-of-origin variance of pairwise distances by a median of about 60%, drew downstream template selections from a smaller and more consistent pool of specimens, and fit held-out specimens more closely in all 62 strains. On the great-ape data the atlases agreed closely in surface geometry, but the downstream morphospace became reference-dependent when the sample was taxonomically imbalanced, and a smaller balanced subset outperformed the larger imbalanced one.
4. Iterative consensus atlas building removes a persistent bias from automated landmarking and yields reference-invariant, reproducible results, with sampling balance mattering more than absolute sample size. Because the atlas stabilises quickly, it can be built from a small balanced subset while the remaining specimens are simply landmarked against it, a practical route to scaling reference-invariant landmarking. The method is implemented in ALPACA within SlicerMorph, with a mock library enabling headless use on HPC.

## Introduction

A common application of image registration is to transfer a set of anatomical references, whether they are segmentations, or coordinates of anatomical landmarks, from a source image to a target one (B. B. Avants et al., 2008; Rohlfing et al., 2004, 2005; Tustison et al., 2014; Young & Maga, 2015). Through this procedure, a large number of similar specimens can be automatically segmented or landmarked and analytical throughput can be increased (Maga et al., 2017).

ALPACA module (Porto et al., 2021) in the SlicerMorph geometric morphometrics toolkit (Rolfe et al., 2021) provides a graphical user interface (GUI) driven way of achieving automated landmarking for 3D models. Its Interactive Mode allows the users to register a pair of models, and transfer the landmarks associated with the source model to the target one while changing the registration parameters and evaluating their effect on the outcome. Once a reliable protocol is established, Batch Mode allows users to process many samples by specifying the folder that contains the models of target specimens, as well as the landmark and model files of the source specimen. In Batch Mode, users can also choose to use more than one source model to estimate the position of landmarks in the target specimen. Because these estimates are in the local coordinate system of the target model, they can be simply averaged or subject to further statistical procedures to implement quality control, to improve estimate the final position of the target landmarks, while reducing bias in the data (Zhang et al., 2022). In both interactive and batch mode the procedure is as follows: dense point clouds (extraction of 4 000–10 000 points are suggested per Porto et al., 2021) are sampled from the target and reference models, Fast Point Feature Histograms (FPFH) descriptors (Rusu et al., 2009) are computed at each point, and a coarse rigid alignment is obtained by FPFH-based RANSAC (Fischler & Bolles, 1981) and refined by Iterative Closest Point (ICP) (Besl & McKay, 1992); bringing the two clouds into a common rigid-body frame (and, with an optional scaling, a common isotropic scale). Non-rigid Coherent Point Drift (CPD) is then applied to deform the aligned source point cloud into the target (Myronenko & Song, 2010). The dense source-to-warped-source displacement field defines a Thin Plate Spline (TPS) transform (Bookstein, 1989), which is finally evaluated at the reference’s hand-placed landmarks to produce the predicted target landmarks. More detailed implementation specific information about ALPACA can be found in both papers (Porto et al., 2021; Zhang et al., 2022).

The unknown and challenging question in this procedure is how to choose the source model(s), particularly for a new and large dataset. If the investigator has done a small pilot, then they probably have enough understanding of the morphological variation in the dataset so that the reference selection process can avoid outliers or individuals that appear to be morphological outliers. This may not always be easy or possible for studies with hundreds to thousands of specimens involved in quantitative genetic or intraspecific variation studies.

The Templates functionality of the ALPACA module was designed to offer an alternative to this bootstrapping challenge (Zhang et al., 2022). The user chooses 3D model arbitrarily to bootstrap the process. This reference sample is downsampled to a sparse point cloud data (PCD) of a few hundred points. Because there are no anatomical landmarks, these PCDs serve as geometrically homologous pseudo-landmarks. Every other specimen is then rigidly aligned to the bootstrap sample via FPFH+RANSAC+ICP, and the bootstrap sample’s PCDs are transferred onto each aligned target by closest-point lookup, yielding a set of pseudo-landmarks that are in correspondence across the sample. These points are then subjected to GPA and PCA to construct a morphospace, and k-means clustering on that morphospace selects the user-specified N specimens nearest each cluster centroid to serve as MALPACA templates.

Unfortunately, the primary vulnerability, not knowing enough about the morphological variation, still applies to this process. Users might unknowingly choose outliers or otherwise morphologically unique specimens as the initial sample and potentially bias the analysis, which subsequently biases the extracted PCDs, the morphospace, and the resulting k-means selection. This cascade has the potential to leave a strong reference-of-origin signal in downstream analyses.

To address this, we propose replacing the arbitrary fixed reference with a consensus atlas built through multiple iterations using a sample of specimens. This consensus atlas building is used commonly in volumetric image registration pipelines (B. Avants & Gee, 2004; Evans et al., 2012; Joshi et al., 2004). By iteratively warping the initial reference toward the average shape of all specimens, the resulting atlas theoretically depends less on the specific starting point. In this paper, we primarily demonstrate and evaluate this iterative consensus model building pipeline using an intraspecific dataset of laboratory mice consisting of wild-derived and classical inbred strains. We test the claims that iterative consensus building is superior to the existing fixed-reference selection, reduces starting-reference influence, and improves out-of-sample fit. Furthermore, we explore the boundaries of this method, discussing the constraints of generalizing this pipeline to highly variable multi-species datasets.

## Materials and Methods

### Mouse Dataset

To assess the improvements associated with the iterative consensus atlas building approach, we utilized a 62-strain *Mus musculus* skull panel of common classical and wild-derived inbred strains (Maga et al., 2017). Each strain is presented by one specimen. This use case represents a very traditional application of the Templates workflow in intraspecies variation studies. Dataset is publicly accessible at https://github.com/SlicerMorpho/Mouse_Models.

### Original Fixed-Reference Pipeline (A)

This is the ALPACA’s Templates behaviour as originally implemented: the bootstrap specimen’s 3D model is downsampled to a sparse matching PCD template (spacingFactor = 0.04, ∼487– 535 points per specimen) and used directly as the template for pseudolandmarks. A GPA and PCA are conducted on these PCDs to create a morphospace, and k-means clustering is used to identify samples closest to the centroids of the specified number of clusters. We set the cluster sizes to 3, 5, 7, and 10.

### New Iterative Consensus Atlas Pipeline (B)

We invoked ALPACA’s new Templates.buildConsensusAtlas method with five iterations and spacingFactor = 0.02. This spacing factor resulted in ∼2,000–2,500 sparse control points per atlas, depending on the bootstrap model. Because the iterative procedure preserves the connectivity graph of the bootstrap 3D model, sutures present in the starting reference would otherwise appear nearly identically in every consensus atlas, creating the false impression that suture shape is unchanged across atlases. The starting reference was therefore Taubin-smoothed (20 iterations, passband 0.1) before iterations began, as the atlas is intended to represent the population’s 3D geometry, not the bootstrap specimen’s suture detail. At each iteration every specimen was similarity-aligned (FPFH+RANSAC+ICP with scaling) to the current atlas; sparse correspondences were obtained by closest-point lookup from the atlas’s downsampled control points onto the aligned specimen and then transported back to the specimen’s original frame by inverting the transform. A partial Procrustes mean of the per-specimen sparse correspondences, anchored to the atlas control-point frame, gave the new target shape. The next atlas was then produced by warping the previous atlas model through a TPS transform defined by the per-control-point displacements between the atlas frame and the Procrustes mean. For this paper, the final model derived at this fifth iteration is considered the “consensus atlas”

The resultant atlas is then used to initialize a sparser PCD extraction (spacingFactor = 0.04, 485-535 pseudo-landmarks). Like the fixed reference pipeline these points are used as the pseudo-landmark template that gets transferred to every specimen in the sample. The pipeline then proceeded through GPA, PCA, and k-means clustering with the same cluster sizes as the fixed-reference pipeline above.

Both pipelines were repeated 62 times, with each specimen serving in turn as the bootstrap reference.

### Statistical Analyses

#### Step-to-step convergence

To verify that the iterative atlas builder converges, we computed the root-mean-square per-vertex displacement between consecutive atlas states (raw model → smoothened → iter00 → iter01 → … → iter04 → consensus_atlas) for each of the 62 reference choices. Because for each atlas, model topology is preserved across iterations, vertices correspond by ID and no nearest-neighbour search is required. However, small rigid-body drift can accumulate from iteration to iteration as each consensus pass re-aligns specimens. To remove that drift before measuring geometric change, every consecutive pair of atlases (A, B) was rigidly aligned in closed form by the Kabsch–Umeyama solution; i.e., the optimal rotation R and translation t minimising ‖R · B + t − A‖^2^ given the known vertex correspondence (Kabsch, 1976, 1978; Umeyama, 1991). Then, the displacement was quantified as step-RMS(A → B) = √(mean_k_ ‖R · b_k + t − a_k‖^2^) over the matched vertex set. This is the single-shot Procrustes equivalent of one ICP step under known correspondence, and it eliminates rigid-body drift from the convergence trace. A monotonically decreasing step-RMS across iterations indicates the atlas is settling into a stable shape.

#### Inter-atlas surface similarity

We also quantified atlas-to-atlas variability directly on the 62 reconstructed atlas models from the new consensus workflow. Because the vertex topology in every consensus atlas is different from each other, we cannot use the approach described above. Instead, the symmetric mean nearest-neighbour (NN) distance was computed on the full models’ vertices after FPFH+RANSAC+ICP alignment was run on subsampled point clouds (∼3,500–5,500 points); the resulting transform was then applied to full-resolution vertices of the source model, and NN distances were queried in both directions against all vertices. For the consensus condition the comparison is between the 62 consensus atlas models; for the fixed-reference condition the corresponding quantity is the pairwise distance between the 62 raw bootstrap models themselves, since the fixed-reference pipeline does not construct a synthetic atlas.

#### Cross-reference morphospace similarity (RV coefficient)

To quantify how similar the downstream morphospaces are across reference choices, we use the RV coefficient (Escoufier, 1973), a basis-invariant multivariate analogue of squared correlation that compares two configurations of the same set of N specimens. For each reference choice i we form the centred Gram matrix 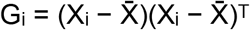 from the N × (p · 3) GPA-aligned PCD configuration matrix X_i_ (where p is the number of sparse landmarks). The RV coefficient between two reference choices is then

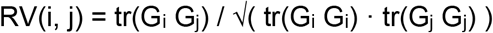

bounded in [0, 1]: RV = 1 means the two morphospaces have identical specimen-by-specimen similarity structure (up to rigid rotation/reflection and overall scale), RV = 0 means no shared structure. We compute RV for all N(N − 1)/2 unordered pairs of reference choices (1891 pairs for the mouse run) and report the mean and minimum as direct measures of how reference-invariant the morphospace is. Higher mean and higher minimum RV indicate that the downstream shape representation is robust to the bootstrap-reference choice. If the iterative consensus atlas building reduces the bias associated with the specimen used to initialize the process, the resulting morphospaces should be similar to each other regardless of the bootstrapping sample.

#### Reference-choice variance of pairwise Procrustes distances

To assess reproducibility of pairwise specimen-shape comparisons across reference choices, we computed the standard deviation (σ) of each pairwise Procrustes distance (PD) across all 62 reference initializations for both the fixed-reference baseline (σ_A) and the consensus pipeline (σ_B). For each of the 1891 specimen pairs we then obtained one paired (σ_A, σ_B) observation. A paired Wilcoxon signed-rank test with the one-sided alternative σ_B < σ_A was applied. If the iterative atlas building reduces reference-of-origin bias, σ_B should be systematically smaller than σ_A and the median σ_B/σ_A ratio should be substantially below 1.

#### Cross-reference Pearson correlations on Centroid Size and Procrustes Distances

As a complementary check, we computed Pearson correlations (Pearson, 1895) between the per-specimen centroid-size (CS) vectors and the per-pair Procrustes-distance (PD) vectors across reference choices, separately under each pipeline. For each pair of reference choices (i, j), we vectorised the 62 CS values under reference i and under reference j and computed Pearson r(CS_i, CS_j); similarly for the 1891 PD values. High mean and minimum r across the N(N − 1)/2 reference pairs indicate that the downstream centroid-size and pairwise-distance vectors are reproducible regardless of which specimen seeded the pipeline.

#### Downstream k-means pick-set stability

To test whether reference invariance propagates into the actual downstream Templates selection workflow, we replicated Templates.templatesSelection exactly (all N − 1 principal components, scipy.cluster.vq.kmeans with thresh = 0, iter = 100, and np.random.seed(1000) set before every call to remove k-means initialisation noise) on each of the 62 reference choices under both pipelines. Each run returned a K-tuple of selected specimens (K ∈ {3, 5, 7, 10}). For each pipeline we measured (a) pairwise Jaccard similarity (Jaccard, 1912) of the K-tuples across the N(N − 1)/2 reference pairs, (b) the number of unique specimens ever picked across the 62 trials, and (c) the Shannon entropy (Shannon, 1948) of the per-specimen pick-frequency distribution. Higher Jaccard means more consistent picks; fewer unique specimens and lower entropy mean the pipeline draws from a tighter pool of MALPACA candidates regardless of which reference seeded the run. A paired Wilcoxon signed-rank test (Wilcoxon, 1945) over the 1891 trial-pair Jaccards tests whether the consensus pipeline is significantly more consistent.

### Modifications to run ALPACA on HPC environments

Due to the large numbers involved, we executed these computational experiments on a high performance computer cluster environment. While 3D Slicer ships with a Python interpreter (SlicerPython) that can be invoked headlessly without launching a GUI, as a GUI driven module, ALPACA’s code paths still depend on certain libraries that are only populated when the full Slicer application is running. To work around this, we wrote a small mock library that stubs out the slicer.* calls the Templates routines invoke. With the mock loaded before importing ALPACA, the relevant Templates logic — matchingPCD, buildConsensusAtlas, and supporting helpers — can be imported and executed under the bare SlicerPython interpreter on HPC nodes without launching the Slicer GUI application. The mock library can be found at https://github.com/muratmaga/slicer-mock. This library is extensible and already supports other SlicerMorph modules (such as GPA), and common Slicer operations like interacting with volumes and markups so that involving them can be run headlessly on HPC environments using the bundled SlicerPython interpreter.

## Results

Across all analyses conducted, all paired comparisons between fixed-reference (A) and the consensus (B) pipelines on the 62-strain mouse dataset returned statistically significant differences in the direction of the consensus pipeline.

### Atlas Convergence and Similarity

The iterative atlas builder converged within five iterations, with mean step-to-step RMS of 0.278 mm in the first iteration and below 0.05 mm from third iteration and onwards (Figure 1).

**Figure 1.**
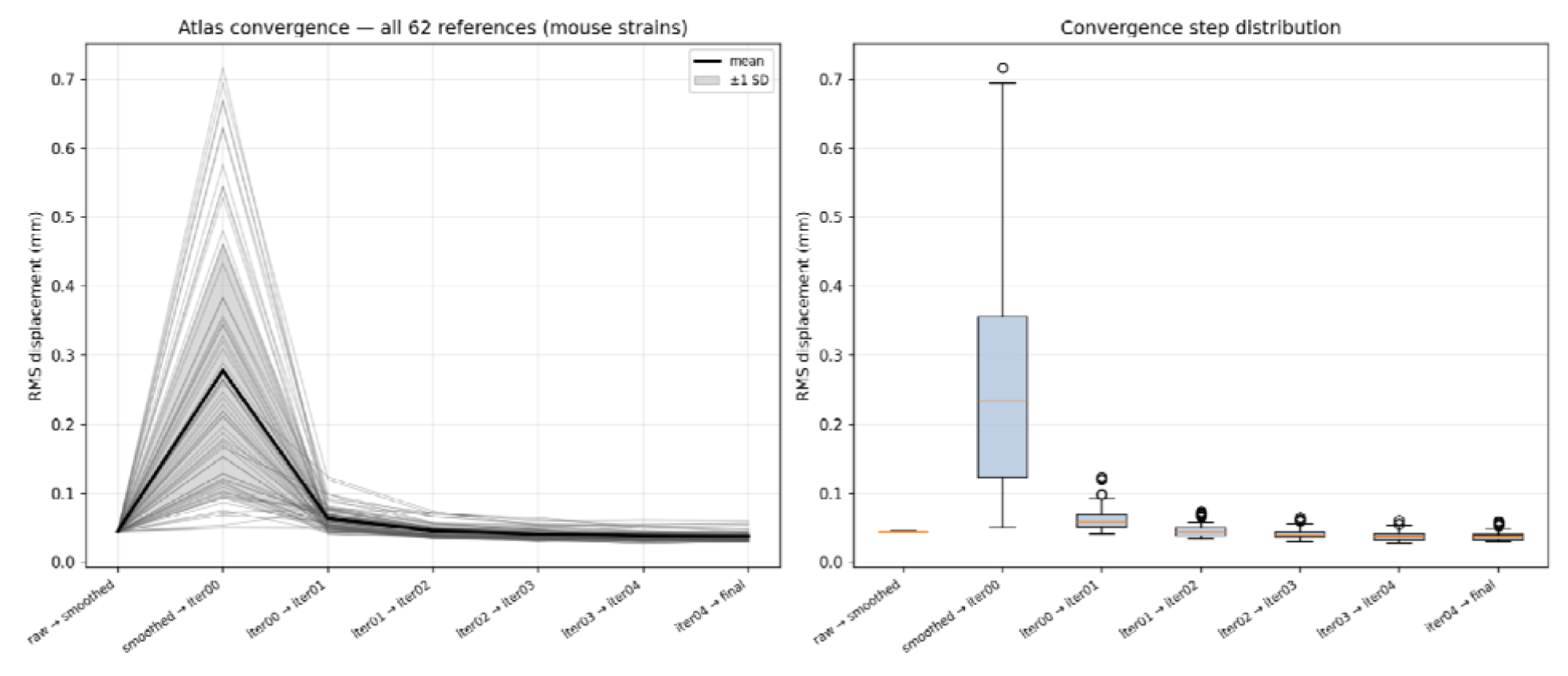
Mouse atlas convergence. **A**. Step-to-step vertex RMS along the bootstrap → smoothed → iter00–04 → final chain, one curve per reference. **B**. Distribution of per-step RMS values across the 62 references, by transition

### Inter-atlas surface similarity

Mean inter-atlas surface distance among the 62 consensus atlases was 0.0835 mm (0.055% of mean centroid size); the corresponding value among the 62 fixed-reference templates was 0.1747 mm (0.116% of mean centroid size).

### Cross-reference morphospace similarity (RV coefficient)

Across 1891 reference pairs, the RV coefficient between PCD-derived Gram matrices was higher under consensus (mean 0.960, SD 0.013) than under fixed reference (mean 0.944, SD 0.019); the paired Wilcoxon signed-rank test gave RV_B > RV_A in 1714 of 1891 reference pairs (W = 56,028, two-sided p = 4.9 × 10^−273^; Figure 2).

**Figure 2.**
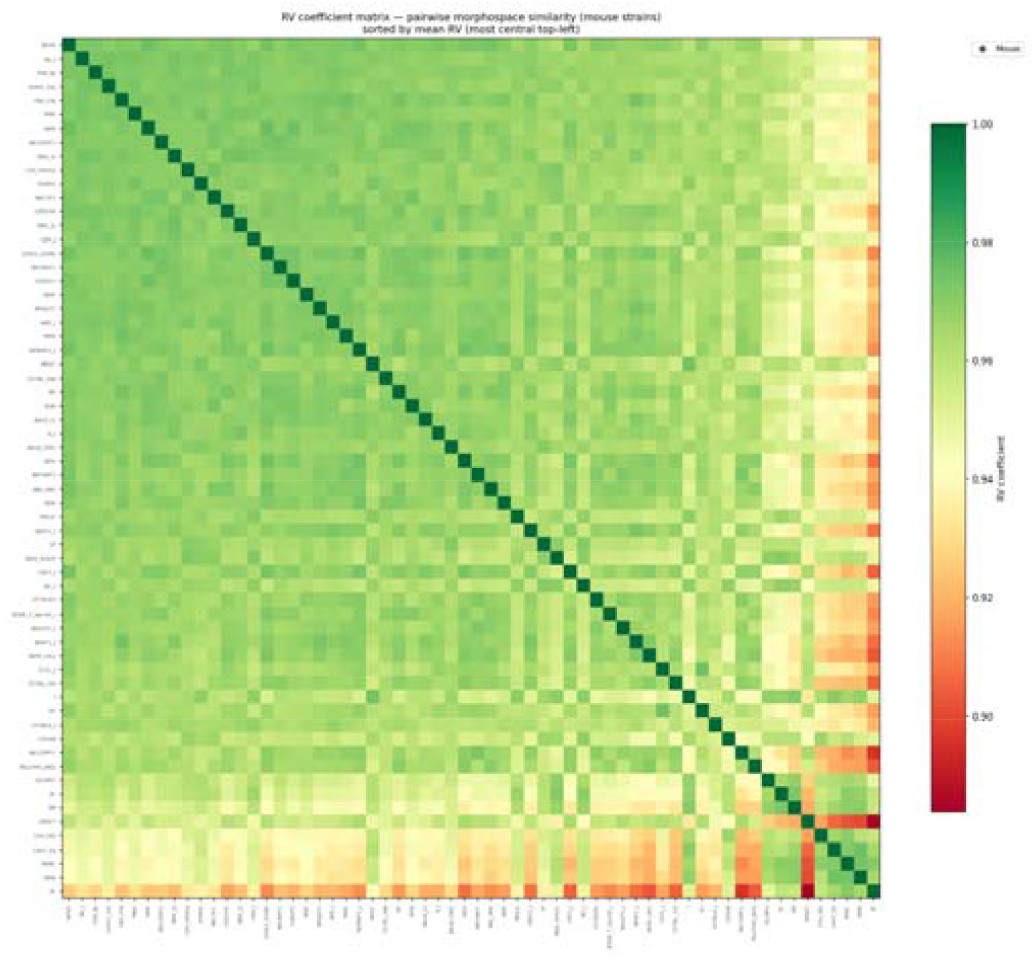
RV coefficient heatmap (mouse, consensus pipeline). Pairwise RV between PCD-derived Gram matrices across the 62 consensus references, ordered by mean centrality (most central top-left).

### Reference-choice variance of pairwise Procrustes distances

For the per-pair reference-choice standard deviation of pairwise Procrustes distance, median σ was 0.094 mm under consensus and 0.245 mm under fixed reference, with σ_B < σ_A in all 1891 specimen pairs (paired Wilcoxon W = 0, one-sided p = 9.4 × 10^−311^); the median difference of 0.152 mm is of the same order as the mean first-iteration atlas displacement (0.278 mm). σ_B/σ_A < 1 for every single pair (Figure 3A). Consensus reduces reference-choice PD variance by a median 60.5% versus fixed reference (Figure 3B). Choosing a different reference shifts the pairwise Procrustes distance between any two mouse strains by a median of ∼245 µm under fixed reference vs ∼94 µm under consensus. For comparison, 245 µm shift is comparable to the difference we observe in the first iteration of atlas building.

**Figure 3.**
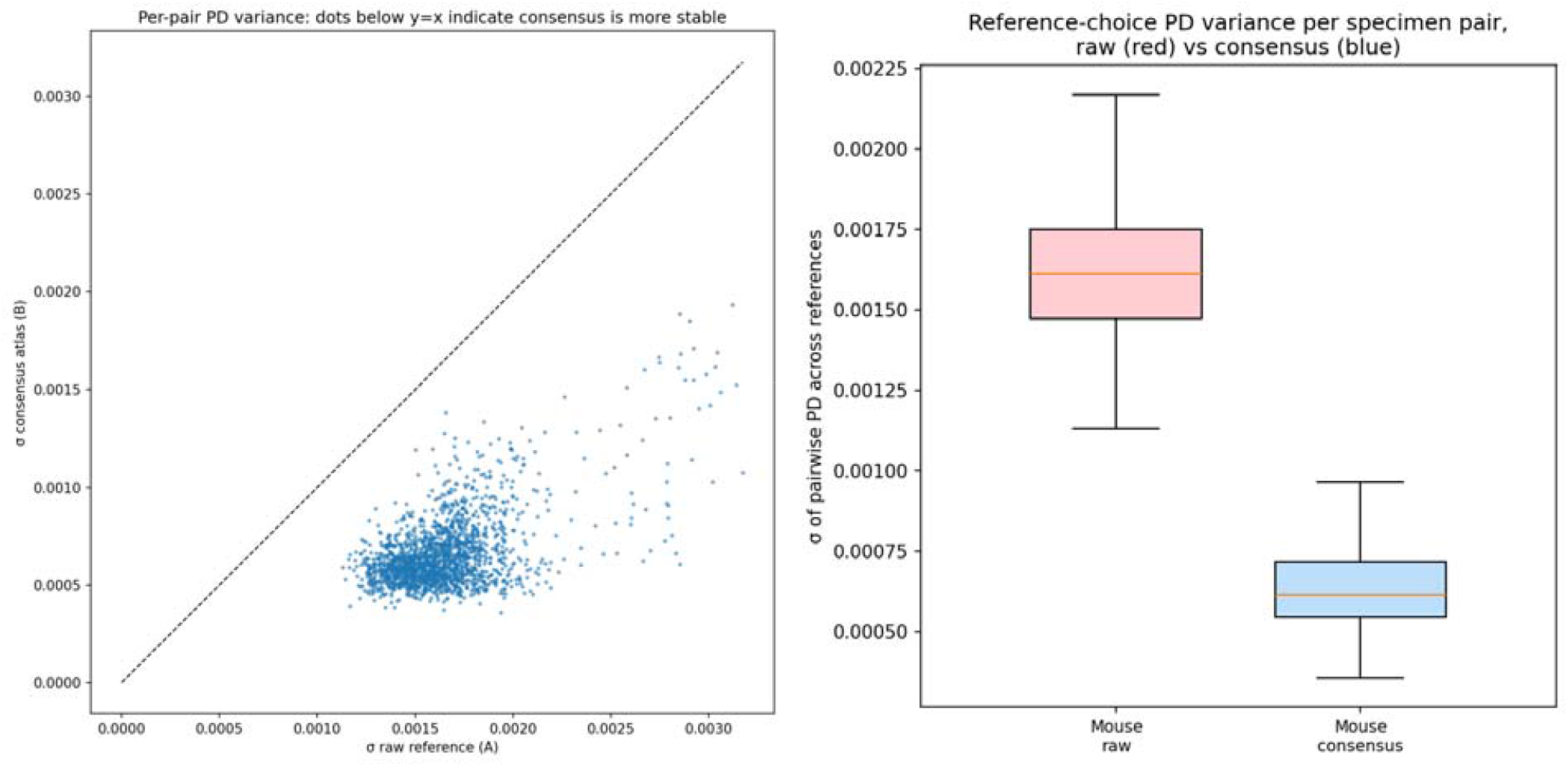
**A. Per-pair PD reference-choice variance, consensus vs fixed reference.** Scatter of σ_B (consensus) against σ_A (fixed reference) over all 1891 specimen pairs; the y = x line marks parity. **3B Distribution of per-pair PD variance, by pipeline**. Boxplot of σ_A and σ_B across all 1891 specimen pairs.

### Cross-reference Pearson correlations (CS and PD vectors)

Cross-reference Pearson correlations on centroid-size vectors were near unity under both pipelines (Figure 4A). Cross-reference Pearson correlations on pairwise-PD vectors were higher under consensus (mean r = 0.960, minimum 0.821) than under fixed reference (mean r = 0.889, minimum 0.579), with B > A in 1889 of 1891 reference pairs (Figure 4B).

**Figure 4.**
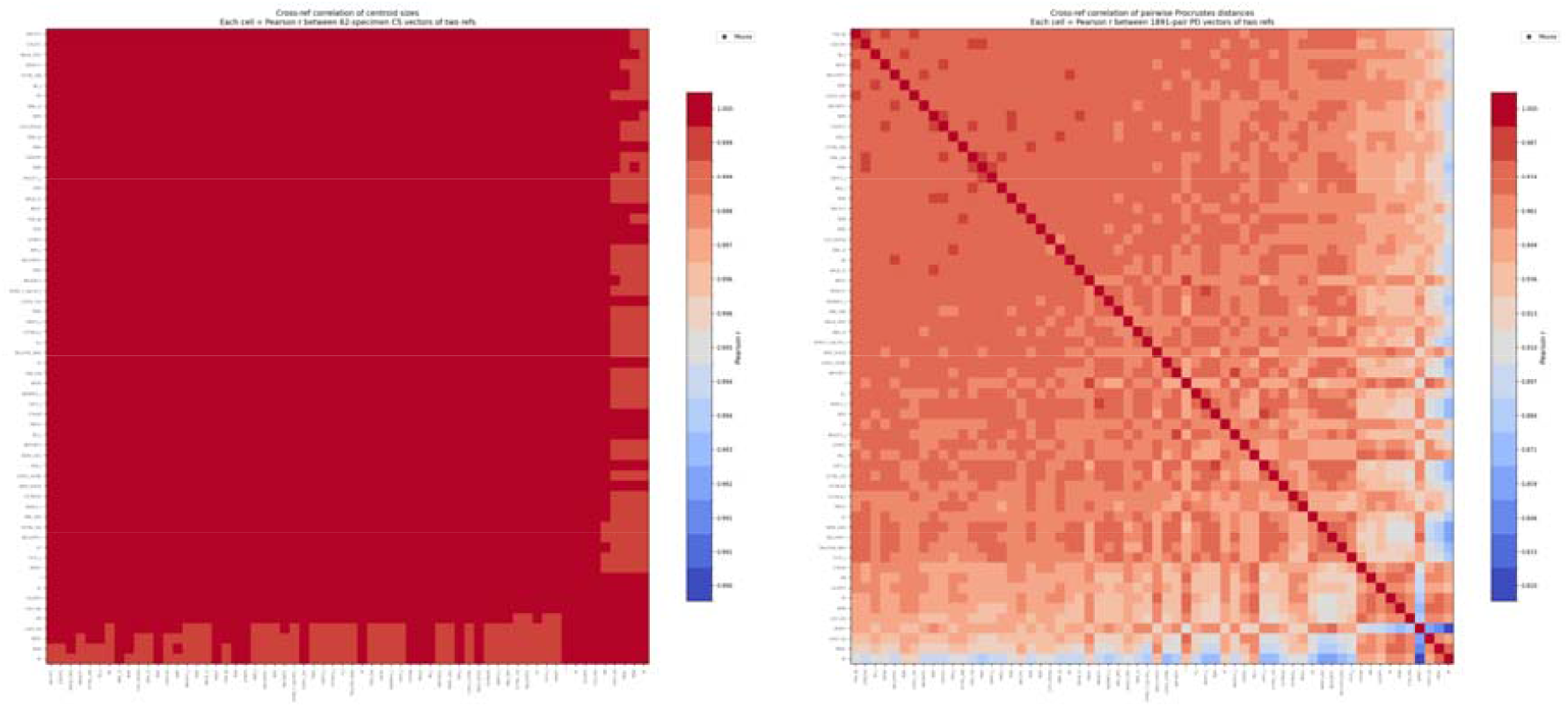
**A. Centroid-size cross-reference correlation for mouse consensus atlas pipeline**. Pairwise Pearson r between per-specimen CS vectors produced under each of the 62 reference choices. **B. Pairwise-PD cross-reference correlation for mouse consensus atlas pipeline**. Pairwise Pearson r between PD vectors produced under each of the 62 reference choices.

### Downstream k-means pick-set stability

The downstream K-means template-selection step drew from a 2-to 4-fold smaller pool of unique specimens under consensus across K ∈ {3, 5, 7, 10}, and mean pairwise Jaccard between K-tuples was higher under consensus at K = 3, 7, and 10 (paired Wilcoxon p ≤ 10^−25^ at each K); the K = 5 case did not reach significance, consistent with the limited Jaccard ladder for 5-element sets (Figure 5).

**Figure 5.**
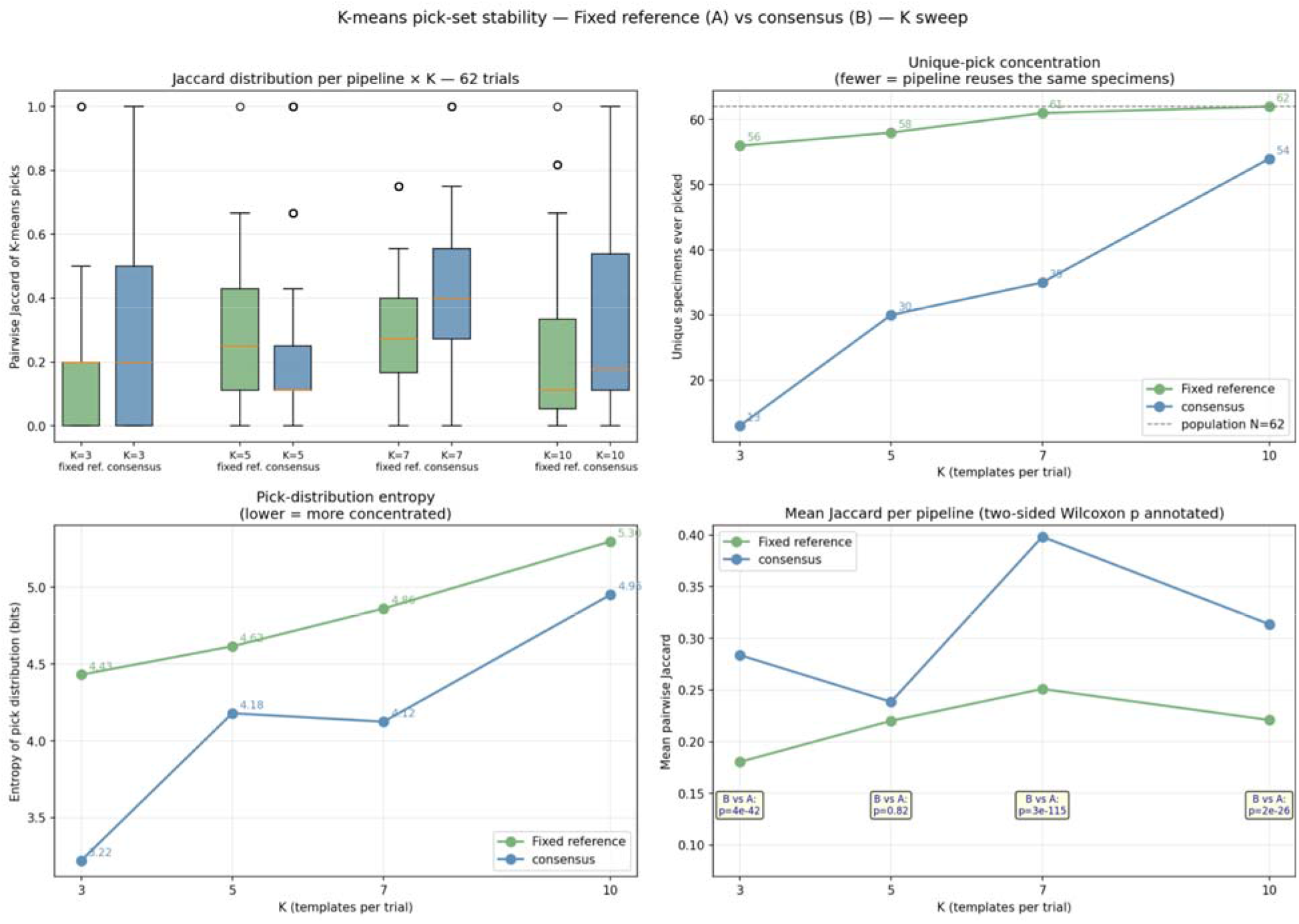
Downstream K-means pick-set stability for mouse pipelines. **Top-left**: per-pair Jaccard distribution. **Top-right**: unique-specimen pool size (fewer = pipeline reuses the same specimens). **Bottom-left**: Shannon entropy of the per-specimen pick-frequency distribution. Bottom-right: mean pairwise Jaccard, with the two-sided paired-Wilcoxon p annotated.

## Discussion

The intraspecific evaluation clearly demonstrates that iterative consensus atlas construction is a vastly superior alternative to fixed-reference selection. By replacing the arbitrary fixed reference with a synthetic mean, we removed a persistent, one-directional bias that propagates throughout the entire automated landmarking pipeline. In the mouse dataset, this translated to a nearly 60% reduction in reference-of-origin noise and highly stable downstream k-means template selections.

A further question is whether this improved internal stability also translates into better predictive accuracy for specimens that did not participate in building the atlas. To test this directly we ran a round-robin leave-one-out experiment. We held out each of the 62 strains in turn. For a given held-out strain, a consensus atlas was rebuilt from the other 61 specimens. The build also depends on which specimen seeds it, so we repeated it once for each of the 61 possible bootstrap seeds. This gives 61 configurations per held-out strain, and 62 × 61 = 3,782 configurations in total. In every configuration the held-out strain was excluded from both the atlas construction and the sample mean it was later compared against. For each configuration we measured out-of-sample fit as the Procrustes distance between the held-out specimen’s pseudo-landmark configuration and the GPA mean of the other 61 specimens. We computed the same quantity for the original fixed-reference Templates pipeline, using the identical held-out design. Because the 3,782 configurations are clustered by held-out specimen and are therefore not independent, we aggregated to one observation per strain, its mean residual across the 61 bootstrap choices, before testing.

The consensus pipeline fit held-out specimens more closely than the fixed-reference pipeline for all 62 strains. The median out-of-sample residual fell from 0.0193 to 0.0176 in dimensionless Procrustes units, a reduction of roughly 8% (paired Wilcoxon W = 0, p = 3.8 × 10^-12). In physical terms, at the mean centroid size of 152.3 mm, this is a drop from about 2.93 mm to 2.68 mm, or close to 0.25 mm. At the level of individual configurations, consensus gave the smaller residual in 3,516 of the 3,782 pairs (93%); we report this as a descriptive count rather than a test, since these configurations are not independent. This 8% reduction in out-of-sample fit is a different kind of result from the 60% reduction in reference-choice variance reported above (Table 4), and the two percentages are not directly comparable. Table 4 is a precision result: it measures how much the distance between two specimens wobbles when the seed reference is swapped, and consensus removes about 60% of that wobble. The leave-one-out test is an accuracy result: it measures how far a held-out specimen lands from the sample mean, averaged over reference choices, and consensus lowers that by about 8%. The two are fractions of very different baselines. The 60% is taken against the template-induced wobble alone, which is itself small (median sigma of 0.0006 to 0.0016 in Table 4). The 8% is taken against the full residual of about 0.018, the bulk of which is the strain’s genuine morphological distance from the average mouse and cannot be changed by any template. In absolute terms the two analyses agree: consensus removes a template effect of roughly 0.001 to 0.0017 dimensionless units, which appears as 60% of the small variance baseline and as 8% of the large residual baseline. The leave-one-out result therefore confirms that the gains from iterative consensus building are not confined to the internal self-consistency of the morphospace documented in Tables 3 to 5, but extend to how well the resulting template generalizes to specimens outside the sample used to construct it.

**Table 1.**
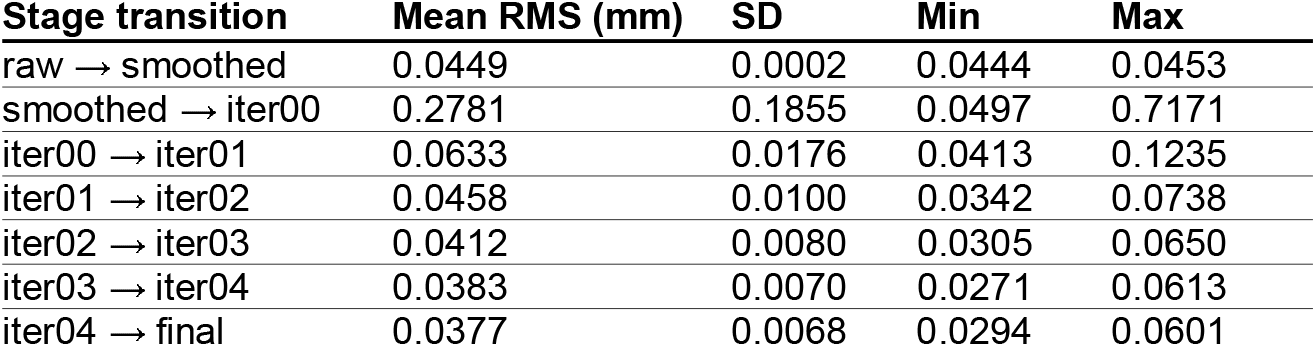
Mean RMS associated with each step of the consensus atlas building.

| Stage transition | Mean RMS (mm) | SD | Min | Max |
| --- | --- | --- | --- | --- |
| raw → smoothed | 0.0449 | 0.0002 | 0.0444 | 0.0453 |
| smoothed → iter00 | 0.2781 | 0.1855 | 0.0497 | 0.7171 |
| iter00 → iter01 | 0.0633 | 0.0176 | 0.0413 | 0.1235 |
| iter01 → iter02 | 0.0458 | 0.0100 | 0.0342 | 0.0738 |
| iter02 → iter03 | 0.0412 | 0.0080 | 0.0305 | 0.0650 |
| iter03 → iter04 | 0.0383 | 0.0070 | 0.0271 | 0.0613 |
| iter04 → final | 0.0377 | 0.0068 | 0.0294 | 0.0601 |

**Table 2.**
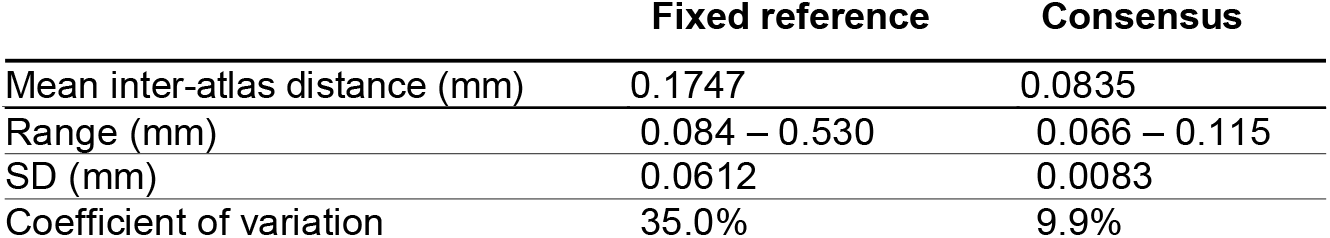
Comparison of inter-atlas difference between fixed reference and consensus atlas pipelines.

**Table 3.** Comparison of morphospaces derived from using a fixed reference and consensus atlas (RV coefficient) and the paired Wilcoxon signed-rank test conducted on 1891 observations:

| Pipeline | Mean | Min | Max | SD |
| --- | --- | --- | --- | --- |
| Fixed reference | 0.944 | 0.872 | 0.981 | 0.019 |
| Consensus | 0.960 | 0.884 | 0.976 | 0.013 |

| Test | Statistic | p (two-sided) | Direction |
| --- | --- | --- | --- |
| Wilcoxon (B vs A) | $W = 56,028$ | $4.9 \times 10^{-2} \square^3$ | $RV\_B > RV\_A$ in 1714 / 1891 pairs (90.6%) |

**Table 4.** Summary statistics and comparison of pairwise Procrustes distance associated with all 62 morphospaces constructed from both fixed reference and consensus atlas pipelines.

| Metric | Median $\sigma_A$<br>(fixed reference) | Median $\sigma_B$<br>(consensus) | Median<br>$\sigma_B/\sigma_A$ | Wilcoxon |
| --- | --- | --- | --- | --- |
| Dimensionless PD | 0.00161 | 0.00061 | 0.395 | $W=0, p = 9.4 \times 10^{-311}$ |
| Absolute (mm) | 0.245 mm | 0.094 mm | 0.395 | (rank-invariant — same W, p) |

**Table 5.**
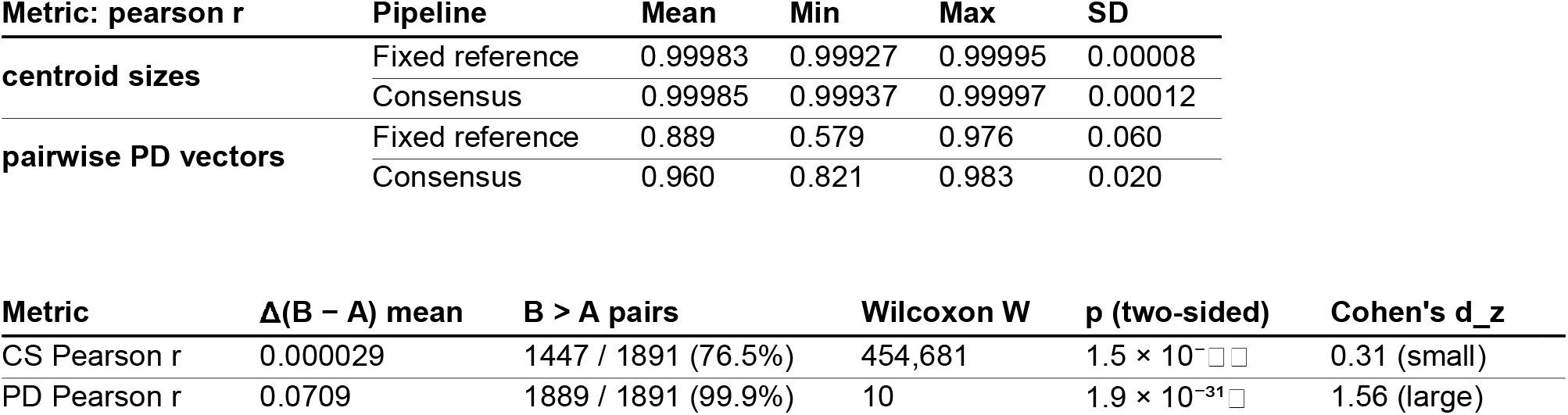
Pearson correlation and Paired Wilcoxon signed-rank tests of centroid sizes and pairwise Procrustes distance associated with all 62 morphospaces constructed.

**Table 6.**
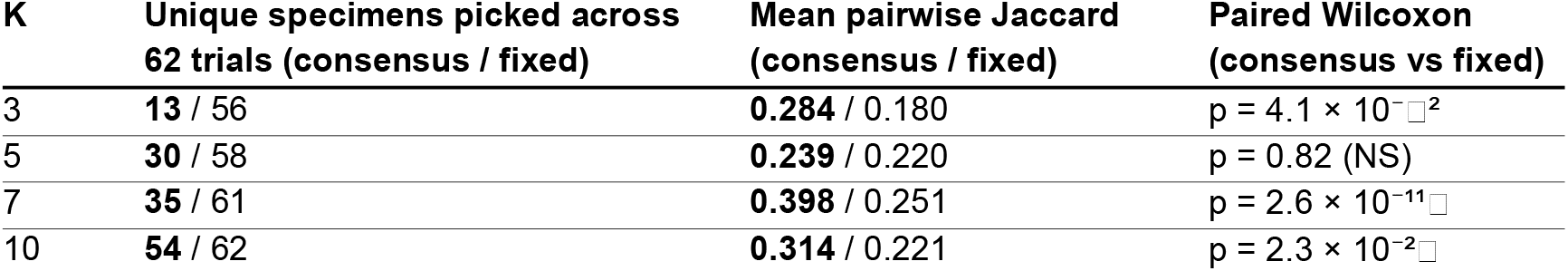
The evaluation of k-mean pick-set stability. Fewer number of unique specimens across all 62 morphospaces indicates more similar morphospaces.

### Generalizing to Multi-Species Problems

The iterative consensus pipeline is effective on canonical, single-species datasets. Whether the same approach generalises to multi-species datasets depends on the morphological variation and sampling design.

To probe these limits we applied the pipeline to an 81-skull great ape dataset (Gorilla, Pan, Pongo), characterised by a 33 % mean centroid-size difference between the largest (Gorilla, mean CS = 1808) and smallest (Pan, mean CS = 1362) species, severe class imbalance (Gorilla n = 38, Pongo n = 30, Pan n = 13), and extreme sexual dimorphism.

Across the 81 reference choices the resulting consensus atlases agreed closely in absolute terms: the mean pairwise inter-atlas distance was 0.72 mm under FPFH+RANSAC+ICP alignment with symmetric nearest-neighbour distance on the full vertex set — about 0.04 % of mean specimen centroid size, comparable in normalised terms to the 0.05 % observed across the 62 mouse atlases. Cross-reference morphospace consistency, however, degraded substantially. The mean pairwise RV coefficient between specimen-coordinate spaces fell from 0.960 in the mouse dataset to 0.729 in the apes, and the cross-reference Procrustes-distance vector correlation fell from near-unity in the mouse case to 0.75 here. Inter-atlas agreement therefore remains tight at the surface level, but the downstream statistical landscape (PCA position, pairwise PD structure) becomes markedly reference-dependent (Figure 6).

**Figure 6.**
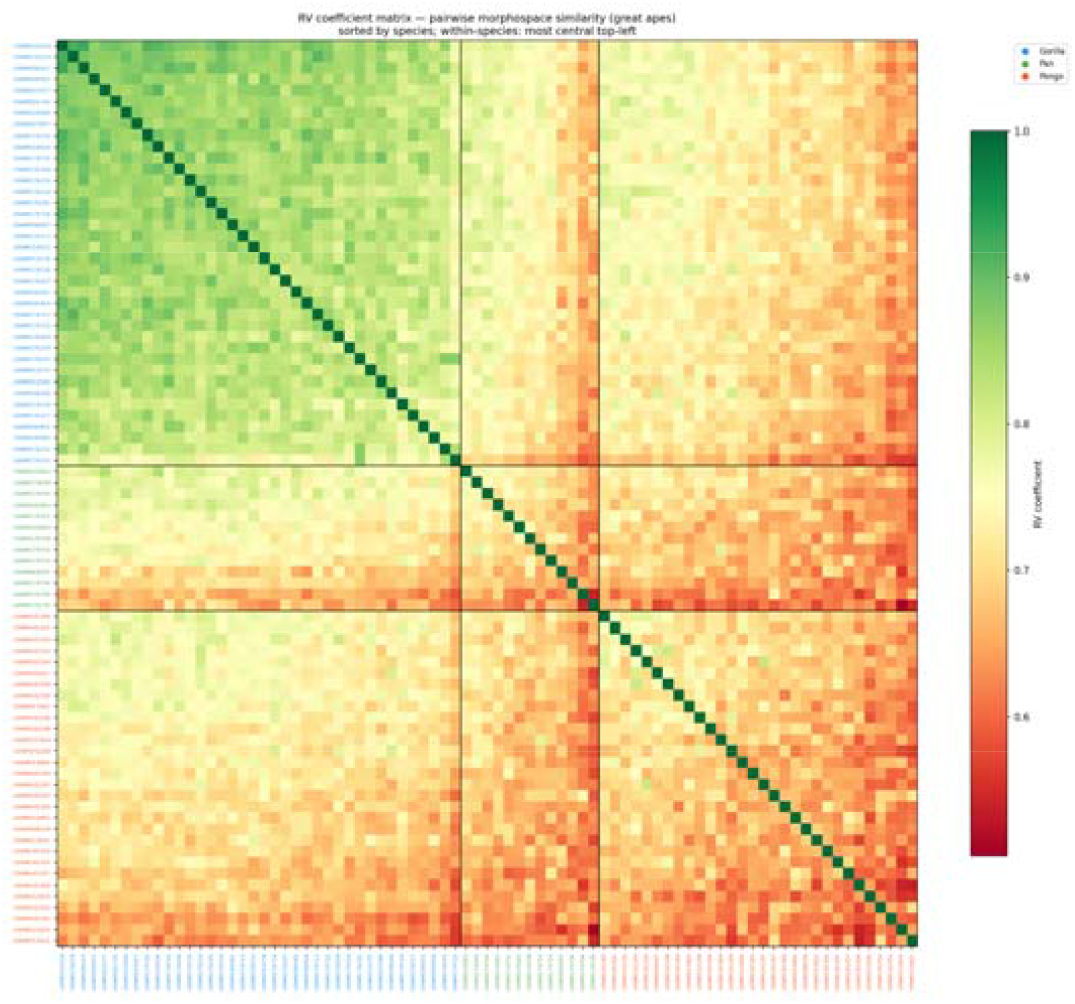
RV heatmap of the apes-full dataset, sorted by species. References sorted by species (Gorilla 38, Pan 13, Pongo 30) and, within each species block, by mean RV to all other references (most central at top-left). The dominant Gorilla block shows tight within-species agreement; the smaller Pan block and the off-diagonal cross-species cells expose the imbalance-driven bias of the consensus morphospace.

### Importance of Sampling Design

Because the consensus is an iterative average, it is dominated by the composition of the sample. In the unbalanced 81-skull dataset the consensus morphospace remained internally consistent for the well-sampled Gorillas but degraded for the under-represented Pan group.

Re-running the pipeline on a balanced subset (MIX: 5 males + 5 females per species, n = 30) produced a substantial increase in cross-reference reproducibility despite the smaller sample (RV mean 0.838 vs 0.729 for the imbalanced 81; PD-vector r mean 0.835 vs 0.751). Conversely, a single-sex design (FEM: 7 females per species, n = 21) showed the weakest pairwise PD reproducibility of the three datasets (mean r = 0.428) and a slightly lower mean RV (0.663) than even the full 81 — indicating that removing the dimorphism axis does not compensate for inadequate within-group sample size. The supplementary figure (Figure S1) shows the consensus atlases produced when a few representative specimens from the study sample were each used to bootstrap the process.

Taken together, these observations indicate that for multi-species or strongly dimorphic targets, sampling balance (equal n per class and per sex) matters more than absolute sample size for reference-invariant outcomes. Where balanced sampling is achievable the iterative consensus approach generalises usefully; where it is not, the downstream reference-dependence of PD-based statistics becomes the dominant source of analytical variability.

We therefore argue that iterative consensus atlas building, as currently implemented, is applicable to multi-species datasets, but only up to a point. Its efficacy and usefulness are governed by between-class size disparity, class imbalance, and within-class dimorphism rather than by absolute sample size, and, perhaps more importantly, by whether the morphology of the included species can be represented canonically in a biologically plausible form. In our great-ape test we expect results would be substantially poorer if the other large ape, humans, were included in the sample: the extreme contrast between the prognathic face of non-human apes and the orthognathic face of humans is unlikely to be reconciled by this fully automated approach without expert guidance, such as dense manual landmarks anchoring the alignment. Conversely, building consensus atlases across multiple families within class Rodentia, with several representatives each, should perform well.

A question that becomes pressing at scale is how many specimens must actually go into building the atlas. The atlas-building sample and the study sample are not the same thing, and they need not be the same size. Every specimen in a study is ultimately landmarked against the atlas, but the atlas itself is a stable mean shape that the iterative procedure reaches well before the full sample is consumed. The leave-one-out experiment on the mouse data showed that a strain left out of construction was represented about as well as one included in it, and the atlas converged and became reference-invariant within a few iterations. For a project of, say, 300 specimens there is therefore no need to run all 300 through the iterative construction. A smaller, deliberately chosen subset will produce an atlas that the remaining specimens fit just as well, and those remaining specimens are then simply landmarked against it.

How that subset is chosen matters far more than how large it is. An atlas is a geometric scaffold for establishing correspondence, not a precise statistical estimate of the population mean, so it needs to span every major source of shape variation rather than to sample any one of them densely. As the balanced thirty-specimen ape design showed, a modest sample stratified across the factors known to drive shape, namely group or species, sex where dimorphism is present, and the range of sizes, will outperform a larger sample drawn without regard to those axes.

We deliberately avoid prescribing a fixed number, because the size a dataset requires depends on how much variation it contains, not on how many specimens it happens to have. A practical procedure follows from the convergence behaviour of the atlas and from our recommendation to pilot any multi-species application. Begin with a small, balanced subset, add specimens stratum by stratum, and inspect the atlas at each iteration to confirm that it remains anatomically plausible. When further specimens no longer move the atlas, the subset is sufficient, and the rest of the sample can be treated purely as targets for landmarking. Used this way, iterative consensus atlas building offers not only a reference-invariant alternative to fixed-template selection, but a practical route to scaling automated landmarking to datasets far larger than the ones examined here.

## Supporting information

Suppmental Figure 1

## ACKNOWLEDGEMENTS

We thank the Smithsonian’s Division of Mammals and Human Origins Program for the scans of USNM specimens used in this research. These scans were acquired through the generous support of the Smithsonian 2.0 Fund and the Smithsonian’s Collections Care and Preservation Fund.

The generative artificial intelligence tool Claude (Opus 4.7; Anthropic) was utilized as a coding assistant to help develop the computational pipeline and implement modifications to the ALPACA source code. The author directed the tool’s use, reviewed and verified all AI-generated code, and took full responsibility for the accuracy, integrity, and functionality of the final software. The agent is acknowledged as a co-author in the repository commits.

## Authors’ Contributions

Murat Maga conceived and executed the study.

## Data availability

Mouse models used are publicly available at https://github.com/SlicerMorph/Mouse_Models. Original 3D scans for great ape skulls should be obtained from the Smithsonian Institution directly. Downsampled version of the reconstructed 3D models of ape skulls used in the study can be found at: https://app.box.com/s/uwfohnl5ia5lxdoaicmzx1ez146q6tzw

