## Supplementary material for "A new reference-invariant consensus template generation method in ALPACA": Suppmental Figure 1

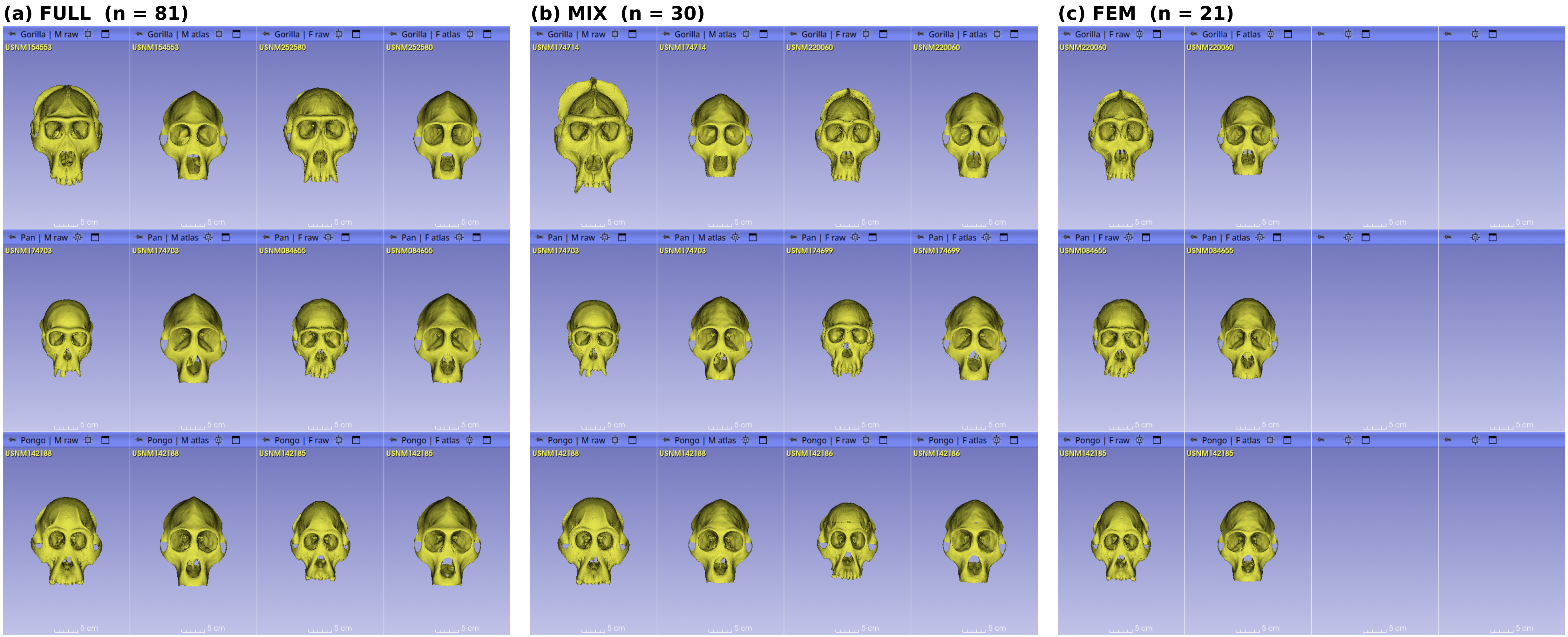


Figure S1. Raw bootstrap specimens and the consensus atlases they produce, across the three great-ape sampling designs. Each panel corresponds to one sampling design: (a) the full imbalanced sample (FULL, n = 81; Gorilla 38, Pan 13, Pongo 30), (b) the balanced mixed-sex subset (MIX, n = 30; 5 males and 5 females per species), and (c) the female-only subset (FEM, n = 21; 7 females per species). Within each panel, rows correspond to species, top to bottom: Gorilla, Pan, Pongo. Columns are organised by sex and representation. For FULL and MIX the four columns are, left to right, male raw, male atlas, female raw, female atlas; for FEM, which contains no males, only the two female columns (female raw, female atlas) are populated and the remaining cells are left blank. In every "raw" column the rendered cranium is an actual specimen from the dataset that was used to bootstrap the pipeline; the adjacent "atlas" column shows the consensus atlas produced when that same specimen seeded the five-iteration consensus build. Each raw specimen and its atlas were rigidly aligned to a common coordinate frame and are shown in anterior view. Despite the substantial morphological differences among the bootstrap specimens in the raw columns (across species, between sexes, and in overall size), the consensus atlases they produce are visually similar to one another, illustrating at the surface level the reference-invariance quantified by the inter-atlas distance analysis.
